# Nucleoli and the nucleoli-centromere association are dynamic during normal development and in cancer

**DOI:** 10.1101/2022.05.21.492846

**Authors:** Aaron Rodrigues, Kyle L. MacQuarrie, Emma Freeman, Kurt Leano, Alexander B Willis, Zhaofa Xu, Angel A Alvarez, Steven Kosak, Yongchao Ma, Bethany E Perez White, Daniel R Foltz, Sui Huang

**Affiliations:** Department of Cell and Developmental Biology, Northwestern University Feinberg School of Medicine. Chicago, IL 60611; Division of Hematology, Oncology, and Stem Cell Transplantation, Department of Pediatrics, Northwestern University Feinberg School of Medicine, Chicago, IL; Department of Biochemistry and Molecular Genetics, Northwestern University Feinberg School of Medicine, Chicago, IL 60611; Departments of Pediatrics, Neurology and Neuroscience, Northwestern University Feinberg School of Medicine, Ann & Robert H. Lurie Children’s Hospital of Chicago, Chicago, IL 60611; Stem Cell Core and Ken & Ruth Davee Department of Neurology, Northwestern University Feinberg School of Medicine. Chicago, IL 60611; Department of Dermatology and Skin Biology and Diseases Resource-based Center, Northwestern University Feinberg School of Medicine. Chicago, IL 60611

## Abstract

Centromeres are known to cluster around nucleoli in drosophila and mammalian cells. However, the functional significance of nucleoli-centromere interaction remains underexplored. We hypothesize that if this conserved interaction is functionally important, it should be dynamic under different physiological and pathological conditions. We examined the nucleolar structure and centromeres at various differentiation stages using cell culture models. The results show dynamic changes of nucleolar number, area, and nucleoli-centromere interactions at differentiation stages and in cancer cells. Embryonic stem cells usually have a single large nucleolus, which associates with a high percentage of centromeres. As cells differentiate into intermediate states, the nucleolar number increases and the association with centromeres decreases. In terminally differentiated cells, including myotubes, neurons and keratinocytes, the number of nucleoli and their association with centromeres are at the lowest. Cancer cells demonstrate the pattern of nucleoli number and nucleoli-centromere association that is akin to proliferative less differentiated cell types, suggesting that nucleolar reorganization and changes in nucleoli-centromere interactions may help facilitate malignant transformation. This idea is supported in a case of pediatric rhabdomyosarcoma, in which induced differentiation inhibits cell proliferation and reduces nucleolar number and centromere association. These findings suggest active roles of nucleolar structure in centromere function and genome organization critical for cellular function in both normal development and cancer.

## Introduction

Nucleoli are multi-functional nuclear organelles beyond being the centers of ribosome synthesis. An increasing number of cellular functions are found associated with this prominent organelle, including signal recognition particle (SRP) assembly, cell cycle regulation, p53 metabolism, stress sensing, gene regulation, and miRNA metabolism [1–3]. More recently, it has been discovered that nucleolar size and area are related to aging in vitro and in vivo [1, 2, 4, 5], further demonstrating the multi-functionality of nucleoli.

Over the past decade, increasing evidence illustrates the spatial interaction of nucleoli with specific domains of chromosomes [3, 6, 7]. Several studies, including next-generation sequencing of nucleolar DNA and HiC experiments demonstrate the association between nucleoli/rDNA and many parts of all chromosomes. These chromosome regions were termed nucleoli associated domains (NADs) [8–12]. NADs are predominantly constitutive and facultative heterochromatin, both in human and mice [13–15]. In general, NAD association with the nucleoli correlates with lower levels of gene expression [3, 15]

The distribution of centromeres is not random, and they are often clustered around nucleoli [16, 17]. In addition, several findings indicate functional connections between centromeres and nucleoli. Centromeres are defined by the presence of a distinct class of nucleosomes containing the centromere-specific histone H3 variant, CENP-A (centromere protein-A). Nucleolar protein NPM1 interacts with CENP-A [18, 19], HJURP, a nucleolar localized CENP-A chaperone [18, 20, 21], and plays a role in centromere assembly. A constitutive centromere-associated network (CCAN) protein, CENP-C, requires an interaction with the nucleolus and RNA for the centromere assembly [22]. Furthermore, in Drosophila, clustering of centromeres around the nucleolus depends on NPM1 and Modulo [23, 24]. These findings underscore the potential functional significance of the nucleoli-centromere association.

To determine the importance of nucleolar-centromere associations in cellular function, we examined the dynamics of these interactions in cell culture models that define multiple distinct developmental stages, including embryonic stem cell, myoblast-myotube, neuroblast-neuron, and epidermal keratinocyte differentiation models. To analyze the pathological redistribution of centromeres, we also compared a panel of normal and cancer cells. Our data suggest that nucleolar structure and nucleoli-centromere interactions are regulated as cells experience different physiological and pathological conditions.

## Results and discussion

### A majority of centromeres associates with nucleoli in pluripotent embryonic stem cells

Embryonic stem cells represent a pluripotent stage of human embryonic development. The H9 embryonic stem cell line was expanded according to established protocols and allowed to differentiate by switching to 10% serum-supplemented DMEM over a 10-day period.

Differentiated cells underwent morphological changes, spreading out and dispersing away from the original stem cell colony. Cells were analyzed by immunofluorescence using antibodies that label nucleoli (Nopp140), centromeres (CREST), and a pluripotent stem cell marker, SSEA4. Cells were imaged through Z sectioning covering entire nuclei and images presented (Fig. 1A) are the maximal projection of the Z stacks to capture all the centromeres within cells. The number of detectable centromeres (individuals with clear separation from one another) and their associations with nucleoli were measured. As shown in Fig. 1A, the undifferentiated H9 cells, generally containing 1-2 large nucleoli, often displayed densely packed nucleoli-associated centromere clusters (Fig. 1A, consider ‘CREST and ‘Merge’ panels). The detected number of centromeres was less than the total number of chromosomes (Fig. 1D) due to their highly clustered nature around the nucleoli, leading to partial overlap of individual centromeres. On average, 71% of centromeres are spatially associated with nucleoli in the undifferentiated H9 cells (Fig. 1E).

**Figure 1.**
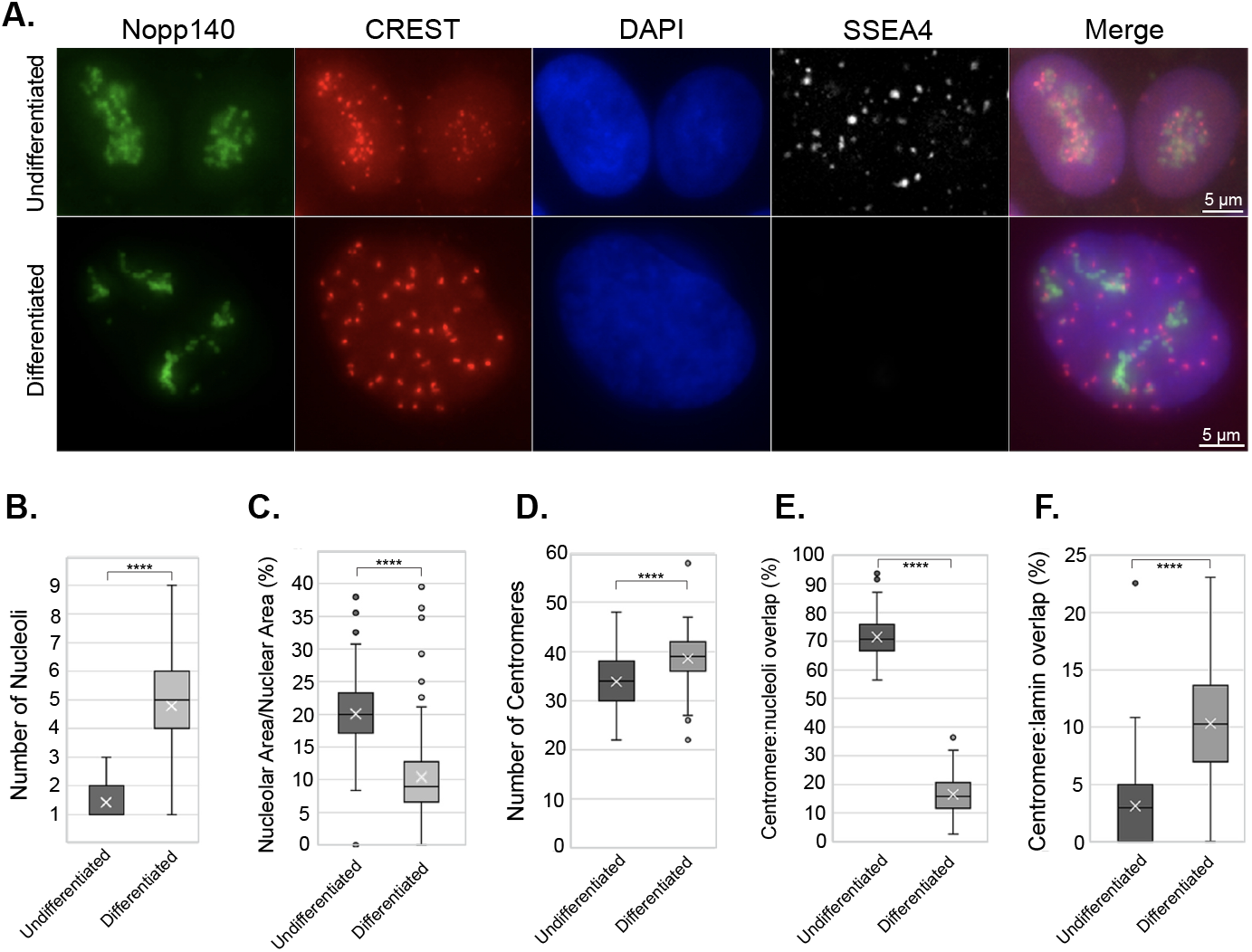
Nucleolus-centromere interactions decrease as H9 human embryonic stem cells differentiate. Nucleoli are immunolabeled with anti-Nopp140 antibody (green) and centromeres by CREST (red) antibodies (A). The pluripotent state of H9 cells was verified through immunofluorescence using SSEA4 antibody, which is lost in differentiating cells (A). Bar=5μm. The number of nucleoli (B) increases as nucleolar area (ratio of nucleolar area over nuclear area) (C). The number of countable centromeres increases as cells differentiate (D). The percentage of centromere associated with nucleoli reaches as high as over 70% in ES H9 cells and reduces significantly as cells differentiate (E). In comparison, percentage of centromeres associating with nuclear periphery is increased (F) **** p<0.0001.

When H9 cells were induced to differentiate, as judged by the loss of the stem marker protein, SSEA4 (Fig. 1A), the cells became more extended, flattened, and spread out. In these cells, the number of nucleoli increased (Fig. 1B), but the association of centromeres with nucleoli was significantly reduced (Fig. 1E). Conversely, the association of centromeres with the nuclear periphery (edge of the DAPI staining) increased (Fig. 1F). These observations demonstrate major nuclear reorganization during ES cell differentiation. Because these cells are proliferating, the significant changes in the nucleolar number may not be due to major fluctuations of ribosome synthesis. This suggests that the reorganization of nucleoli and centromere association correspond to changes of cellular function such as gene expression regulation during differentiation that require specific topological nucleoli and genomic interactions.

### Nucleoli-centromere association is reduced as cells differentiate towards the terminally differentiated state

To evaluate subsequent stages of differentiation, from blasts to terminally differentiated cells, we utilized three types of differentiation models: myoblasts to myotubes, neuroblasts to neurons, and 3D human skin equivalent tissue development.

Primary human myoblasts were induced to differentiate into myotubes. Cells were similarly prepared and immunolabeled as for the ES cells. To ensure cells are fully differentiated in the myotube analysis, only multi-nucleated cells were assessed (Fig. S1). As myoblasts differentiate into myotubes, there is a significant reduction in the number of nucleoli (Fig. 2A and B), nucleolar area (Fig. 2C), and nucleoli-associated centromeres (Fig. 2A and E), while no significant change was noted in association of centromeres with the nuclear periphery (Fig. 2F). The evaluation of nucleoli and centromere localization showed a reduction of total number of countable centromeres (Fig. 2A and D). Such a reduction could be explained by the fusion of centromeres or by a reduction in CENP protein retention in differentiated myotubes as previously reported [25]. Consistent with the latter, examination of publicly available expression data comparing myoblasts and myotubes demonstrated significant downregulation of multiple genes related to centromere structure and function (Table S1) [26]. These findings demonstrate a dynamic reorganization of nucleolar structure and nucleoli-centromere association as cells progress through the myo-differentiation process.

**Figure 2.**
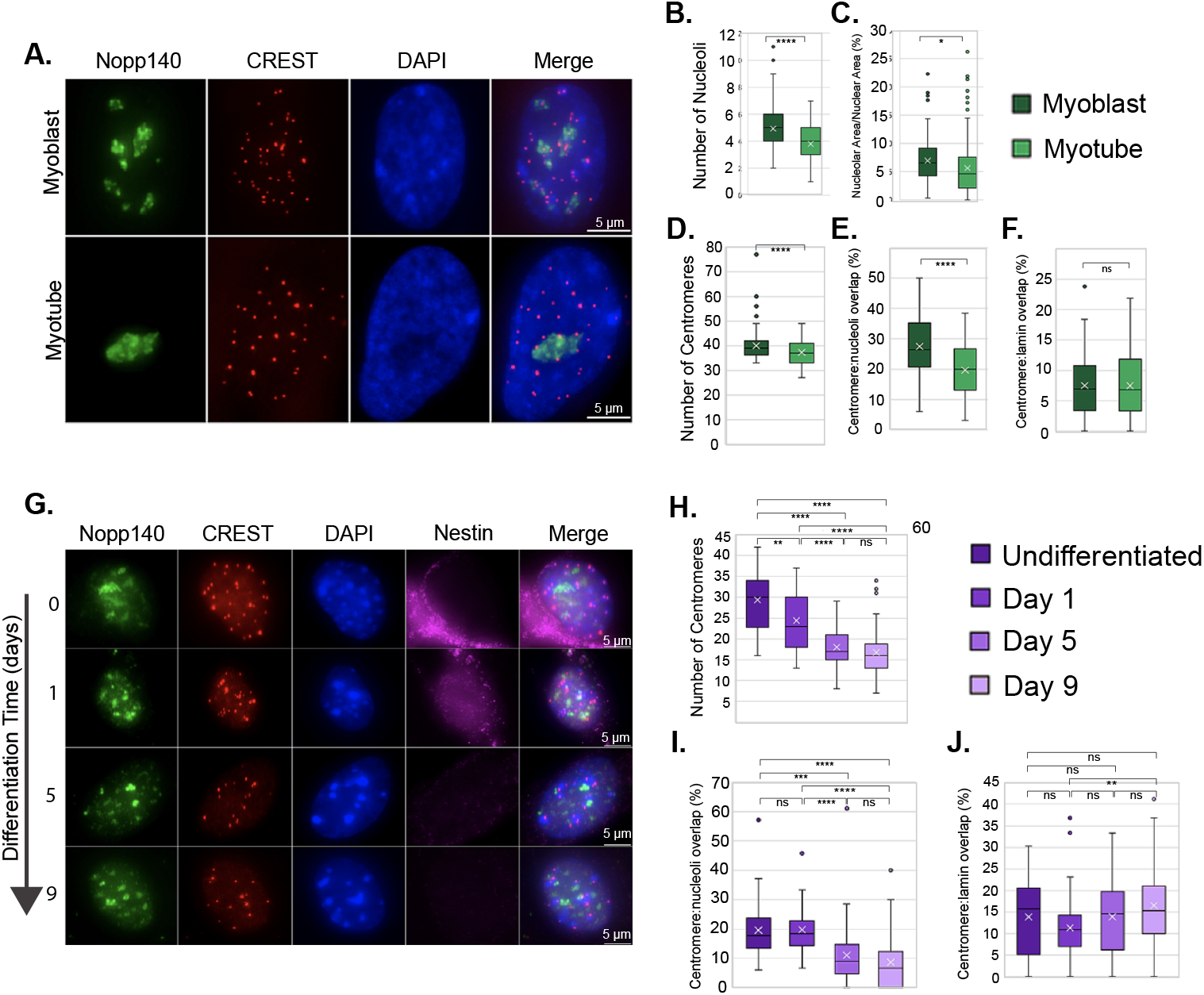
Nucleolus-centromere interactions decrease as myoblasts or neuroblasts differentiate into myotubes or neurons. Nucleoli are immunolabeled with anti-Nopp140 antibody (green) and centromeres by CREST (red) antibodies (A and G). Bar=5μm. Only the nuclei in myotubes (more than one nucleus in a cell) are considered differentiated myocytes. The number of nucleoli (B) and nucleolar area (C) decreases as myoblasts differentiate into myotubes. In addition, the number of countable centromeres (D) and the percentage of centromeres associated with nucleoli (E) reduce significantly as cells differentiate into myotubes. However, the percentage of centromeres associating with nuclear periphery is unchanged (F). Nestin (purple) is a marker of undifferentiated neuroblasts. The number of countable centromeres (H) and the percentage of centromeres associated with nucleoli (I) reduces as cells differentiate into neurons. However, the percentage of centromeres associated with nuclear periphery is unchanged (J). **** p<0.0001.

The second system utilizes the differentiation of mouse neuroblasts to neurons. Neuroblasts were isolated directly from mouse brain and were induced to differentiate. Cells were analyzed at day 0 (neuroblast), 1, 5, and 9 to examine changes throughout differentiation. Nestin expression was used as a marker for undifferentiated neuroblasts. As cells differentiate, judged by the loss of nestin signals (Fig. 2G), the number of detectable centromeres and the percentage of nucleolar-associated centromeres reduces significantly (Fig. 2H and I). Significant changes in centromeres’ association with the nuclear periphery were not observed (Fig. 2J).

To further confirm the dynamics of nucleoli-centromere interactions through differentiation, we employed 3D human skin equivalents (3D HSE) derived from primary human keratinocytes. In this model, the morphogenesis and differentiation closely recapitulate that of normal human epidermis [27, 28]. In addition, mature (day 12) 3D HSE provide a unique advantage because a single tissue cross-section encompasses cells at all stages of epidermal differentiation. This allows for an unbiased view of nucleoli and centromeres characteristics through the differentiation process within a single image. In the basal layer, where epidermal stem cells reside, keratinocytes proliferate. Cells exit the cell cycle and move from the basal layer into the spinous, then the granular layer, where the cells are increasingly differentiated. Finally, cells terminally differentiate and undergo specialized cell death and enucleation in the cornified layer. This process of differentiation is initiated by culture at the air-liquid interface and occurs over the course of up to 12 days (Fig. S2). As shown in Fig. 3A, the immunolabeling of a 6-day 3D HSE culture demonstrated clearly definable layers of cells at different stages of epidermal differentiation. The number of nucleoli (Fig. 3C) and nucleolar area (Fig. 3D) decrease as cellular differentiation progresses, except for layer 2 where the nucleolar area increased compared to the basal layer. It is interesting to note that fibrillarin (Fig. 3A, green signal, arrows), a nucleolar pre-ribosomal processing factor, displays a significant cytoplasmic presence as cells depart from the basal layer and exit the cell cycle, suggesting potential alterations of nucleolar function as cells no longer divide from this layer on.

**Figure 3.**
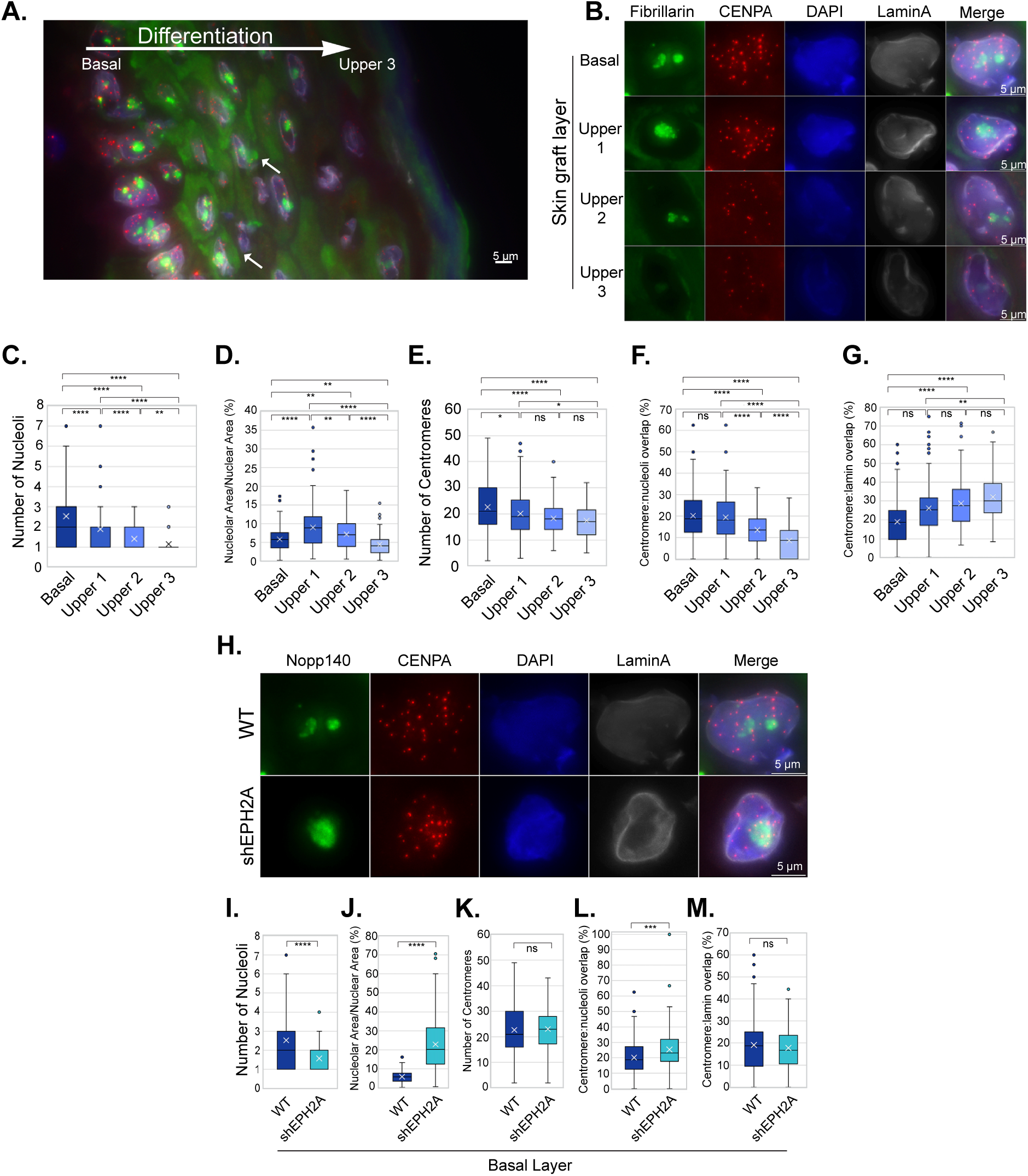
Nucleolus-centromere interactions decrease as karatinocytes differentiate from basal cells towards cornified layers. Nucleoli are immunolabeled with anti-fibrillarin antibody (green), centromeres by anti-CENPA (red) antibodies, and nuclear lamin by anti-lamin A (light purple) antibodies. (A) A cross section of a skin graft containing all layers in the same image demonstrates the clear changes in centromere numbers and intensity of the signals as well as their association with nucleoli as they develop into cornified layers. The large white arrow indicates the direction of differentiation, and small white arrows indicate examples of extranuclear fibrillarin staining. (B) Typical results of stains in cells from layers ranging from basal to upper layers are as shown. Bar=5μm. The number of nucleoli (C) and nucleolar area (D) decrease in parallel with the differentiation. The number of countable centromeres (E) and the percentage of centromeres associated with nucleoli (F) also reduces as cells differentiate. Correspondingly, the percentage of centromeres associating with nuclear periphery is increased (G). (H) Representative stains in basal layer keratinocytes that either were transduced with a short hairpin construct targeting EPH2A (shEPH2A) and therefore with impaired differentiation or with a construct with a scrambled sequence with no target. Keratinocytes with EPH2A knockdown (shEPH2A) demonstrate lower number of nucleoli (I), an increase in nucleolar area relative to nuclear area (J), no difference in centromere number (K), an increase in overlap between centromeres and nucleoli (L) and no difference in centromere overlap with the nuclear periphery (M). **** p<0.0001.

CENP-A antibody staining shows that the number of centromeres (Fig. 3B and E) and nucleoli-centromere interactions decrease (Fig. 3B and F) as keratinocytes differentiated towards the cornified layer. Interestingly, the percentage of centromeres associated with nuclear periphery increases (Fig. 3B and G). When all the differentiation layers are viewed on the same section, it is notable that the labeling signal of CENP-A (red) is significantly reduced as cells differentiate (Fig. 3A). The reduction of centromere numbers is observed across all three terminal differentiation systems and is consistent with the findings in myoblast differentiation where CENP-A deposition to centromeres were reduced [25] in addition to reduction in CENP gene expressions (Table S1).

To further evaluate the relationship between nucleoli-centromere association and differentiation, we asked whether this association changes when differentiation is inhibited. EPHA2, a receptor tyrosine kinase, plays an important role in epidermal differentiation and cancer prevention [29–32]. To block keratinocyte differentiation, we knocked down EPHA2 in primary human keratinocytes using lentiviral shRNA and generated 3D HSE. Compared to the control culture, the EPHA2 depleted keratinocytes show a lack of differentiation and stratification (Fig. 3H). Nucleolar number reduces (Fig. 3I) but nucleolar area increases (Fig. 3J). While there is no significant difference in the centromere number between control and EPHA2-deficient 3D HSE in the basal layer (Fig. 3H and K), the nucleoli-centromere association increases when EPHA2 is depleted (Fig. 3H and L), while the association of centromeres with the nuclear periphery does not change (Fig. 3M). These data further corroborate the idea that differentiation couples with the reduction of nucleoli-centromere interactions.

### Nucleoli-centromere interactions increase when cells are transformed into immortalized or cancerous cells

To further evaluate the dynamics of nucleoli-centromere interactions during differentiation, we examined a cohort of human cancer cells and a normal cell line, as a key feature of cancer is the disruption of differentiated characteristics of the normal tissues of origin [33]. If nucleoli-centromeres interaction decreases as cell differentiate, we would expect the interaction to increase in cancer cells. We examined nucleoli and centromere dynamics in non-cancerous primary human umbilical vein endothelial cells (HUVEC), hTERT-immortalized retinal pigment epithelial cells (RPE1^−hTERT^), colorectal adenocarcinoma cells (DLD-1), metastatic cervical adenocarcinoma (HeLa), and osteosarcoma cells (U2O2). The results (Fig. 4A) show that the number of nucleoli and nucleolar areas varied among the cancer cells, with cancer cells trending towards having more nucleoli and increased area relative to normal cells (Fig. 4B and C). The number of centromeres (Fig. 4D) is significantly increased in all three cancer cell lines compared to diploid cells, consistent with established aneuploidy in these cell lines. The percentage of nucleoli-centromeres increases in immortalized cells (Fig. 4E), and increases further in cancer cells (Fig. 4E). In addition, the percentage of nuclear periphery associated centromeres decreases significantly in hTERT-immortalized and transformed cancer cells (Fig. 4F). These observations indicate that carcinogenesis induced dedifferentiation coincides the increases in nucleoli number, area and nucleoli-associated centromeres and decreases in nuclear periphery association.

**Figure 4.**
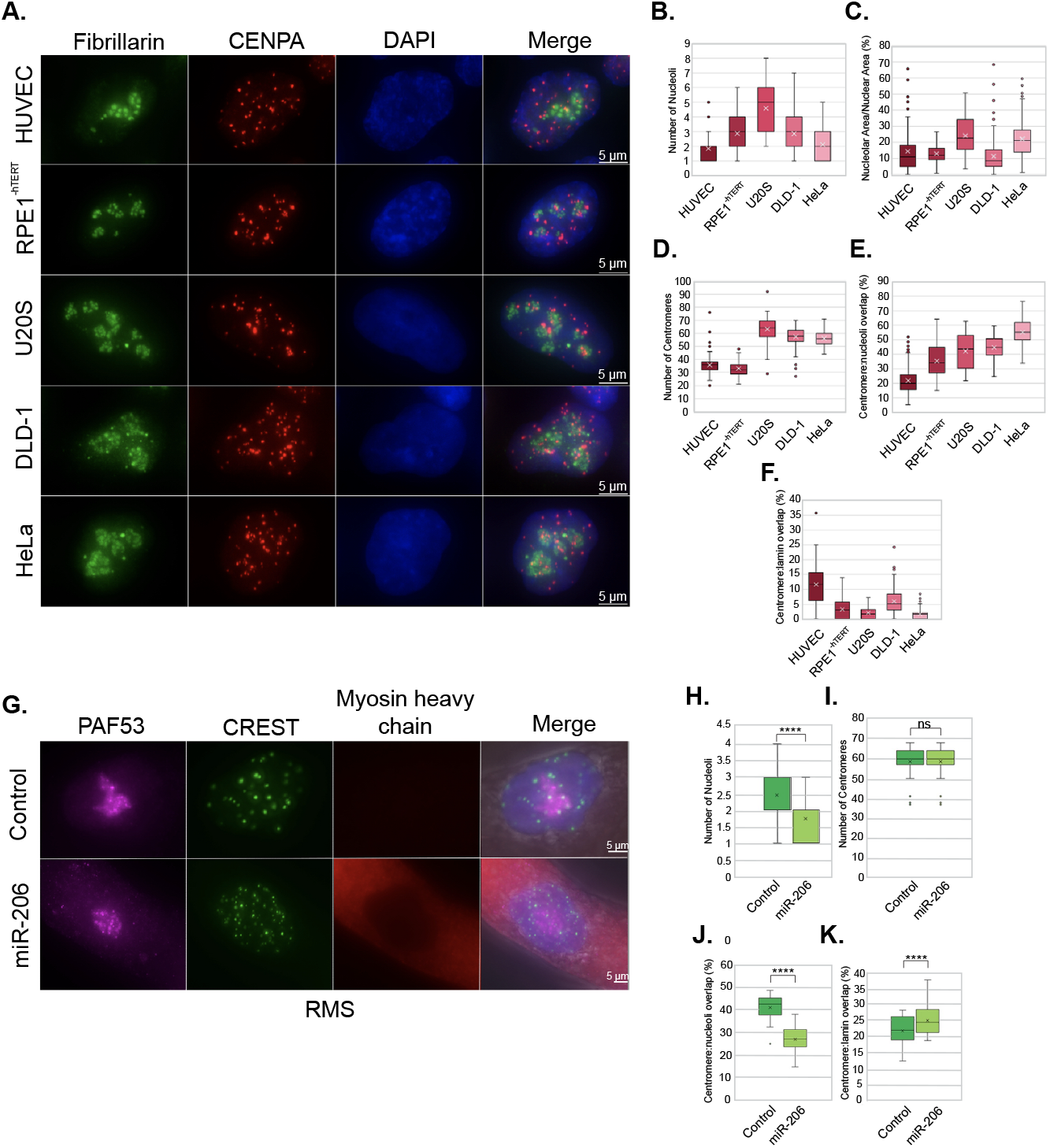
Nucleolus-centromere interactions increase in cancer cells compared to normal cells and reduce upon differentiation. Nucleoli were immunolabeled either with anti-fibrillarin antibody (green) and centromeres by anti-CenpA (red) antibodies (A) or anti-PAF53 (cyan) and CREST (green) (G). Compared to normal HUVEC cells, various immortalized and cancer cells show increases in the number (B) and area of nucleoli (C). While the number of centromeres vary greatly in cancer cells (D), the percentage of nucleolar associated centromeres increases significantly compared to the normal control (E). Conversely, a decreased spatial association of centromeres to the nuclear lamin is seen (F). When pediatric rhabdomyosarcoma cells were induced to differentiate upon treatment with miR-206 (G), the number of nucleoli (H) was reduced, while the number of centromeres did not change (I). Differentiated cells demonstrated a significant reduction in nucleolus-centromere overlap (J), along with a significant increase in centromere-nuclear periphery overlap (K). **** p<0.0001. Bar=5μm.

To further confirm that the dynamics of nucleolar structure and nucleoli-centromere association parallels the differentiation process, we asked whether induction of differentiation in a cancer cell line would restore the state of nucleolar structure and the nucleoli-centromere association in differentiated cells. RD cells are a cell culture model derived from a pediatric rhabdomyosarcoma (RMS). Increased expression of the pro-myogenic miRNA miR-206 has been demonstrated to induce differentiation of RMS cells and inhibit RMS tumor growth in vivo [34, 35]. We examined and compared RMS cells transiently transfected with a miR-206 mimetic or a negative control mimetic and then kept in myogenic differentiation media for 48 hours. The differentiation of these tumor cells into a more myotube-like state was determined by the expression of sarcomeric myosin heavy chain, a structural protein upregulated during normal myogenic differentiation [36] (Fig. 4G). The number of nucleoli was reduced upon differentiation (Fig. G and H). Furthermore, the nucleoli-centromere association (Fig. 4G and J), but not the number of centromeres (Fig. 4G and I) significantly decreased in myosin expressing cells. These data are consistent with our observations of the reduction of nucleoli-centromere association in terminally differentiated cells. In line with what was observed in normal myoblasts and myotubes, staining for specific CENP proteins demonstrated decreased CENPT signal specifically in myosin heavy chain positive RD cells, though CENPA staining appeared unaffected (Fig. S3). Examination of publicly available expression data comparing RD cells to myotubes also demonstrated multiple centromere-related genes expressed at significantly higher levels in the RD cells compared to myotubes (Table S2).

In summary, we report the dynamic nature of nucleolar number, area, and nucleoli-centromere interactions throughout cellular differentiation in several cell types. Nucleolar number and area do not always correspond to each other. The average number of nucleoli in ES cells is 1.5, but the nucleolar area is the largest with a nucleoli/nucleus area ratio of 20%, whereas in blast stage, while nucleolar numbers increased to 3-5 in human cell models, the nucleolar area decreased. In terminally differentiated human cells nucleolar number and areas decrease further. These findings demonstrate that the nucleolar number and area are unlikely to be related to each other or to ribosome synthesis in cycling cells, suggesting roles for nucleolar organization in cellular differentiation.

The nucleoli-centromere association is the highest in pluripotent stem cells and lowest in the terminally differentiated state, in which possibly only the NOR containing chromosome centromeres still associated with nucleoli. The association is directly linked to the differentiation state as either the block of differentiation in a 3D HSE graft by shRNA against an essential epidermal differentiation factor EPHA2 or the dedifferentiation during carcinogenesis significantly increases the nucleoli-centromere association. Furthermore, when a rhabdomyosarcoma cell line was induced to differentiate by the pro-myogenic miRNA miR-206, the association decreases as cells increased expression of myosin heavy chain, a marker of myogenic differentiation, and exit cell cycle [34].

Centromeres are special areas of chromosomes that enable equal segregation of chromosome into the resulting daughter cells during cell division. Centromeres are composed with highly repetitive DNA sequences occupied by centromere specific CENP-A nucleosomes flanked by histone H3 containing nucleosomes that form the peri-centric heterochromatin [37]. The spatial link of heterochromatin with nucleoli is considered a regulatory mechanism that assists the suppression of gene expression [38]. From pluripotent to terminally differentiated cells, heterochromatin is found surrounding nucleoli [38]. ES cells generally have the least amount of heterochromatin [39, 40] (Fig. 5). As the association of centromeres with nucleoli is the highest in ES cells, it is difficult to imagine that the changes of nucleoli-centromere interactions through differentiation stages are simply due to spatial shifting of bulk heterochromatin (Fig. 5). Rather, this suggests a model in which the shifting nucleoli-centromere association reflects a differentiation state-specific role for centromere function and/or genome organization in stem cells. Taken together with our data in other model systems, our findings implicate the importance of nucleolar structure and its interaction with centromeres throughout both differentiation stages and during tumorigenesis

**Figure 5.**
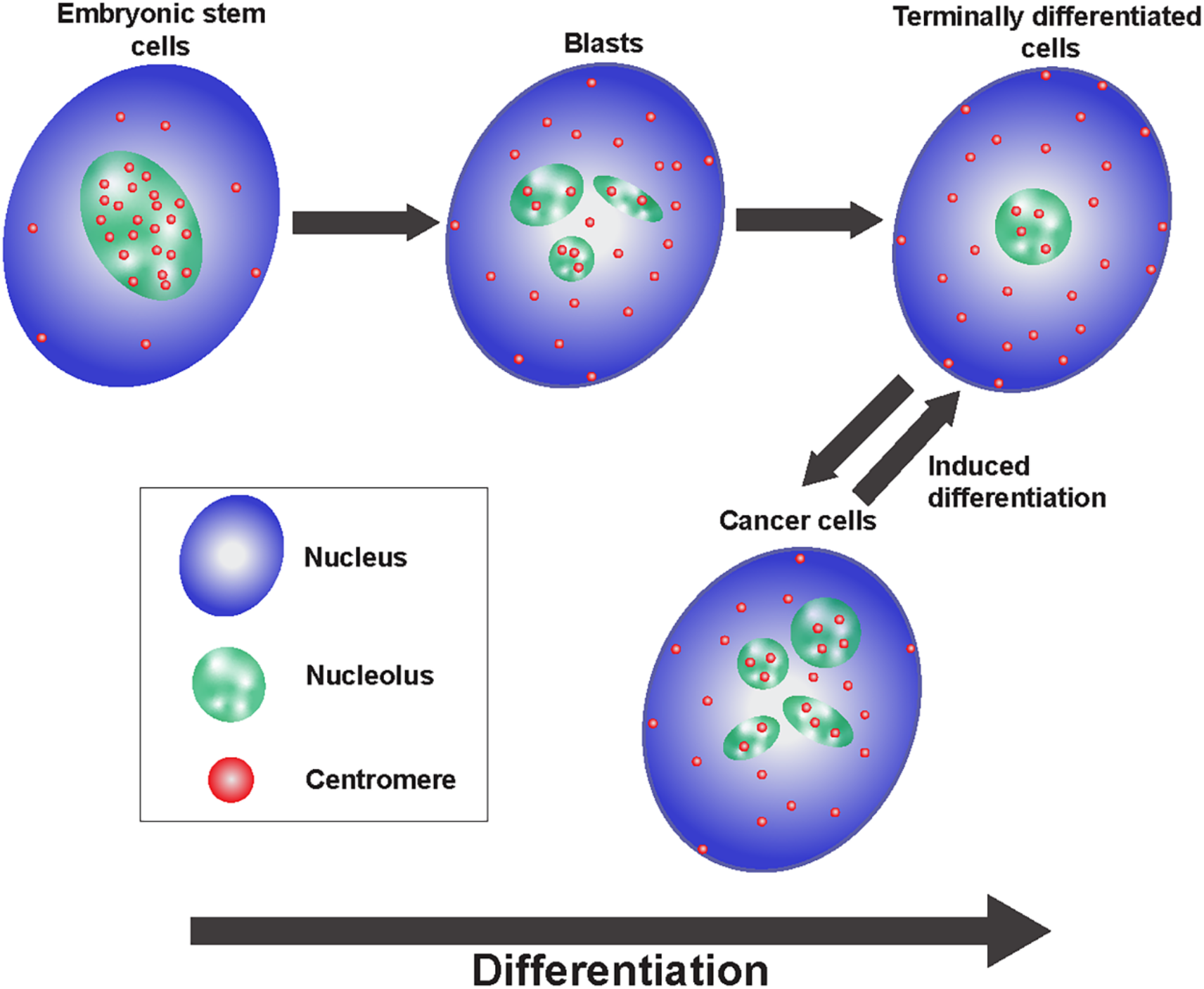
Schematic model demonstrating that the association of centromeres with nucleoli decreases as cell differentiate, is increased in cancer cells, and decreases once more as cancer cells are induced to differentiate.

## Material and Methods

### Immunostaining

The cells were fixed with 4% (weight/volume) paraformaldehyde in PBS for 15 minutes, washed with PBS, solubilized in 0.5% (volume/ volume) Triton X-100 (Sigma) in PBS for 10 minutes, washed, incubated in human CREST antibody at a dilution of 1:10, mouse myosin heavy chain MF-20 antibody at 1:100, or rabbit POLRIE antibody at 1:100 (Proteintech cat no: 16145-1-AP) for 1 hour, washed, and finally incubated in anti-human, anti-mouse, and anti-rabbit Alexa-fluor secondary antibodies at 1:200 (Invitrogen) for 1 hour. The cells were then mounted using Vectashield antifade mounting media with DAPI (Vector Laboratories, Inc. H-1200).

### Embryonic stem cells growth and differentiation

The NIH-approved and registered embryonic stem cell line NIHhESC-09-0022 (H9 ESCs) were acquired from WiCell and expanded using plates or glass coverslips coated with hESC-qualitied Matrigel (Corning) in serum-free mTeSR stem cell medium (Stem Cell Technologies). ESCs were characterized based on morphology to grow in flat and tightly-packed colonies with well-defined borders comprised of cells with high nuclear-to-cytoplasmic ratios and the ability to expand and differentiate. Cells were certified to have a normal karyotype and free from contaminants. ESCs were differentiated by switching to 10% serum-supplemented DMEM (Corning) over a 10-day period.

### 3D Human Skin Equivalents (HSE)

3D HSE were established as previously described using primary neonatal foreskin-derived keratinocytes [27]. Cells were transduced with a lentiviral construct containing a short hairpin sequence targeting EPHA2 or a scrambled sequence (pLKO.EV and pLKO.shEPHA2, gifts of Bingcheng Wang, Case Western Reserve University, Cincinnati, OH) as previously described [32].

### Myoblast differentiation

Primary human skeletal muscle cells were acquired from ATCC and cultured in Mesenchymal Stem Cell Basal Medium (ATCC) supplemented with the Primary Skeletal Muscle Cell Growth Kit (ATCC) components (FBS, dexamethasone, L-glutamine, EGF, FGF-b, and insulin) per manufacturer’s recommendations, as well as 1% penicillin-streptomycin (Gibco). Cells were kept at low density (<50% confluency) during propagation, and 0.25% Trypsin-EDTA (Hyclone) used when passaging cells.

For myoblast samples, cells were split onto untreated coverslips at a low density, cultured in growth media for 16 hours, and were then washed with phosphate-buffered saline (PBS) twice prior to being fixed in 4% paraformaldehyde in PBS for 10 minutes at room temperature. Before fixation, all myoblast samples were inspected to ensure confluency was <50%. For differentiated myotube samples, cells were split at a sufficient density onto untreated coverslips to reach 100% confluency after 16 hours in growth media, then washed three times with PBS before being changed to Skeletal Muscle Differentiation Tool media and cultured for a further 96 hours. Cells were then fixed as above for myoblast samples.

### Rhabdomyosarcoma cells

The RD human rhabdomyosarcoma cell culture line was acquired from ATCC and cultured in DMEM (Gibco) with 10% Fetal Bovine Serum (Hyclone) and 1% penicillin-streptomycin (Gibco). One day prior to transfection, cells were trypsinized (Hyclone) and plated onto untreated glass coverslips (VWR) at sufficient density to reach 50-60% confluency the following day. Cells were transfected using either a miR-206 miRNA mimetic (Thermofisher) or negative control mimetic #1 (Thermofisher) using RNAiMax (Thermofisher). Transfections were performed according to manufacturer’s protocol with the following modifications: the mimetic’s final concentration was 16 nM, no antibiotics were present in the media, and 1.5 uL of RNAiMax was used per well for a 12 well dish. After 24 hours, cells were washed 2x with PBS, and media was changed to low serum differentiation media (DMEM +1% horse serum (Hyclone) + 1x insulin-transferrin-selenium (Corning) + 1% penicillin-streptomycin). Cells were fixed after 48 hours in differentiation media as noted above for myoblasts.

### Neuroblast differentiation

Neuroblasts/neural precursor cells are isolated and grown as described [41]. Dorsal forebrains from timed-pregnant E13.5 mouse embryos were digested with Accutase (Fisher). Neuroblasts were carried on plates coated with Matrigel (Corning) at 80ug/ml and maintained in DMEM-F12 medium (GIBCO) supplemented with B27 (GIBCO), N2 (GIBCO), and Glutamax (GIBCO). A growth factor cocktail containing EGF (epidermal growth factor) (20ng/ml final) (PeproTech) and basic FGF (fibroblast growth factor) (20ng/ml) (PeproTech) in Heparin (5ug/ml) was added to the medium fresh. Cells were carried at densities not exceeding 80%, and all experiments were performed on density- and passage-matched cultures. Cells were incubated in standard conditions: 37°C with 5% CO_2_.

#### Differentiation

Neuroblasts were seeded onto coverslips or tissue culture dishes coated with 20ug/ml poly-L-Lysine (PLL) and 4ug/ml Laminin in NSC medium without growth factors (DMEM-F12 supplemented with N2, B27, Glutamax without bFGF and EGF). For 24-well plates, 300,000 neuroblasts were seeded into each well. The following day, day *in vitro* 1 (Div1), cells were changed into fresh medium containing 50ng/ml BDNF (brain derived neurotrophic factor), 25ng/ml GDNF (glial cell derived neurotrophic factor) and 10uM Forskolin. Half of the culture media in each well were replaced with fresh medium containing BDNF, GDNF, and Forskolin every three days until cells are fully differentiated into neurons.

## Acknowledgements and Funding

D.R.F. is funded by GM1111907 and U10CA260699. Y.C.M. was supported by NIH R01NS094564 and R21NS106307. B.E.P.W. was supported by NIH NIAMS grants K01AR072773 and P30AR075049. S.H. was supported by U10CA260699. K.L.M. is supported, in part, by the National Institutes of Health’s National Center for Advancing Translational Sciences, Grant Number KL2TR001424, as well as the Hyundai Hope on Wheels Scholar Hope Grant, the Stanley Manne Children’s Research Institute, and the Ann & Robert H. Lurie Children’s Hospital of Chicago. The content is solely the responsibility of the authors and does not necessarily represent the official views of the National Institutes of Health. We would also like to thank Drs. Robert Goldman for the CREST and lamin antibodies, Thomas Meier for Nopp140 antibodies, and Joe Ibarra for technical assistance. We would also like to thank Drs. Stephen Adam, Paul Kaufman, and Anastassiia Vertii for critical reading.

