## Supplemental Figures and Tables for "Nucleoli and the nucleoli-centromere association are dynamic during normal development and in cancer"

**Supplemental Materials**:

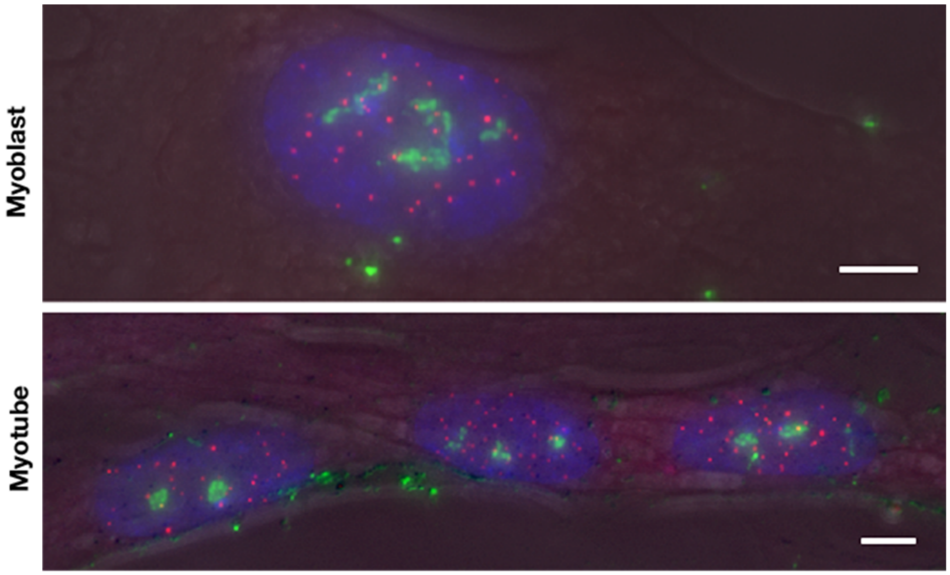


**Supplemental Figure 1.** Differentiated myotubes are those form multi-nuclei tubes. Nucleoli are immunolabeled with anti-Nopp140 antibody (green signals), and centromeres are immunolabeled with anti-CENPA antibody (Red). Only cells in the myotubes were counted as terminally differentiated cells. Bar=0.5 mm

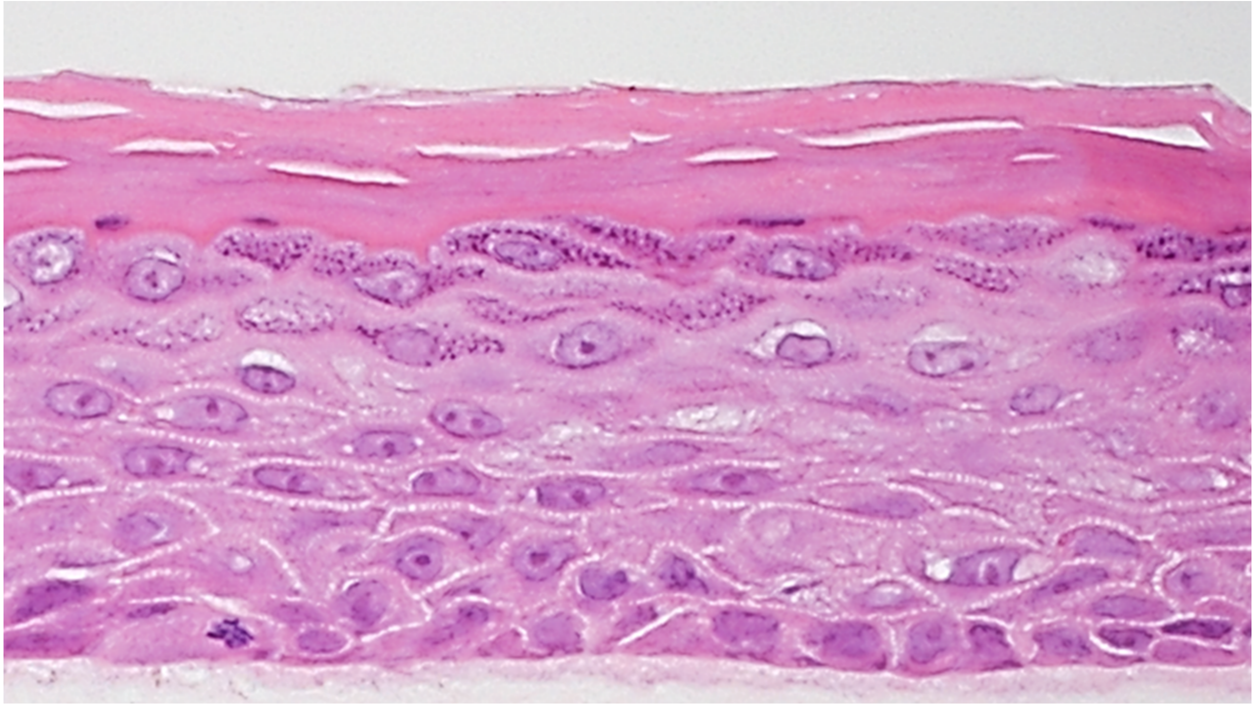

**Supplemental Figure 2.** A representative image of a H&E stained 3D human skin equivalents derived from primary human keratinocytes.

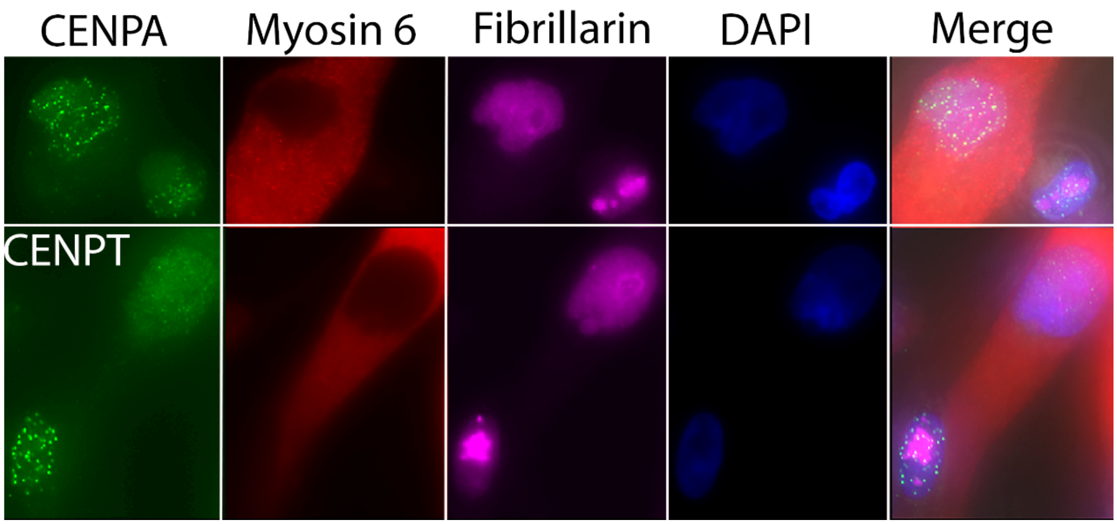


**Supplemental Figure 3.** While CENPA labeling (green, top row) does not appear reduced when cells become differentiated (myosin heavy chain expressing cells, red), CENPT labeling (green, bottom row) significantly reduces in myosin heavy chain expressing cells (red).

Supplemental tables:

| Entrez ID | Gene Symbol | Gene Name | Myoblast:Myotube fold difference (log2 value) |
|---|---|---|---|
| 79019 | CENPM | centromere protein M | -3.16 |
| 55355 | HJURP | Holliday junction recognition protein | -3.07 |
| 1058 | CENPA | Centromere protein A | -2.45 |
| 387103 | CENPW | Centromere protein W | -2.35 |
| 55839 | CENPN | Centromere protein N | -2.31 |
| 11339 | OIP5 | Opa interacting factor 5 | -2.14 |
| 1062 | CENPE | Centromere protein E | -1.96 |
| 2305 | FOXM1 | Forkhead box M1 | -1.87 |
| 55320 | MIS18BP1 | MIS18 binding protein 1 | -1.83 |
| 1063 | CENPF | Centromere protein F, 350/400 kDA | -1.77 |
| 64105 | CENPK | Centromere protein K | -1.54 |
| 1870 | E2F2 | E2F transcription factor 2 | -1.10 |
| 3619 | INCENP | Inner centromere protein antigens | -1.10 |
| 54069 | MIS18A | MIS18 kinetochore protein homolog A | -1.09 |
| 201161 | CENPV | Centromere protein V | -1.07 |

**Supplemental Table 1.** Centromere-related genes significantly downregulated with normal myogenesis.

| Entrez ID | Gene Symbol | Gene Name | RD:Myotube fold difference (log2 value) |
|---|---|---|---|
| 1870 | E2F2 | E2F transcription factor 2 | -3.51 |
| 55355 | HJURP | Holliday junction recognition protein | -3.46 |
| 79019 | CENPM | centromere protein M | -3.11 |
| 11339 | OIP5 | Opa interacting factor 5 | -2.71 |
| 1058 | CENPA | Centromere protein A | -2.54 |
| 55839 | CENPN | Centromere protein N | -2.35 |
| 1063 | CENPF | Centromere protein F, 350/400 kDA | -2.20 |
| 2305 | FOXM1 | Forkhead box M1 | -2.11 |
| 1062 | CENPE | Centromere protein E | -1.88 |
| 201161 | CENPV | Centromere protein V | -1.74 |

**Supplemental Table 2.** Centromere-related genes significantly upregulated in RD rhabdomyosarcoma cells compared to normal myotubes.
